# Damaged mitochondria recruit the effector NEMO to activate NF-κB signaling

**DOI:** 10.1101/2022.06.21.496850

**Authors:** Olivia Harding, Erika L.F. Holzbaur

**Author notes:** Correspondence: Erika L.F. Holzbaur, University of Pennsylvania Perelman School of Medicine 638A Clinical Research Building, 415 Curie Boulevard, Philadelphia, PA 19104. Open access data: https://doi.org/10.5281/zenodo.6654160. Provisional protocols are available through Protocols.io. https://www.protocols.io/view/hela-culture-transfection-and-labeling-of-halo-fus-14egn7rdmv5d/v1. https://www.protocols.io/view/fixation-and-imaging-of-hela-cells-after-mitochond-cbrdsm26. https://www.protocols.io/view/live-imaging-to-investigate-mitophagy-kinetics-and-cbrfsm3n. https://www.protocols.io/view/rna-extraction-and-quantitative-pcr-to-assay-infla-cbrism4e. https://www.protocols.io/view/image-processing-to-investigate-nemo-recruitment-a-cbrjsm4n. https://www.protocols.io/view/cell-lysis-and-gel-electrophoresis-for-protein-ana-cbrksm4w.

## Abstract

Failure to clear damaged mitochondria via mitophagy disrupts physiological function and may initiate damage signaling via inflammatory cascades. However, signaling mechanisms leading from impaired mitophagy to neuro-inflammation are unclear. We discovered that NF-κB essential regulator NEMO is recruited to damaged mitochondria in a Parkin- and p62/SQSTM1-dependent manner in a time-course similar to recruitment of the structurally-related mitophagy receptor, OPTN. NEMO and p62 colocalize, partitioning into distinct domains from OPTN. Either depletion of p62 or mutation of NEMO’s ubiquitin-binding domain abolishes NEMO recruitment, indicating multifactorial interactions. The active catalytic IKK component phospho-IKKβ colocalizes with NEMO on damaged mitochondria, initiating NF-κB signaling and the upregulation of inflammatory cytokines. These findings suggest that damaged mitochondria serve as an intracellular platform for innate immune signaling by promoting the formation of activated IKK complexes in a Parkin-dependent manner. We propose that mitophagy and NF-κB signaling are competing pathways regulating the response to cellular stress.

## Introduction

Selective autophagy is a mechanism by which damaged organelles or macromolecular complexes are targeted for lysosomal clearance (Stolz, Ernst, and Dikic 2014). Specific receptors are recruited via “eat-me” signals, with ubiquitination being one of the most universal (Goodall, Kraus, and Harper 2022; Pohl and Dikic 2019; Rose et al. 2016). Targets as diverse as insoluble protein aggregates, cytosolic *Salmonella*, and depolarized mitochondria are ubiquitinated before they are sequestered and degraded in a conserved process that likely evolved as a mechanism to keep cells safe from harmful waste and invasive microorganisms (Manifava and Ktistakis 2022; Wild et al. 2011). Mutations that disrupt selective autophagy have the potential to delay clearance of damaging entities, and thus may trigger downstream stress pathways such as inflammation.

In the mitophagy pathway for the selective autophagy of damaged mitochondria, receptors are recruited to the outer mitochondrial membrane (OMM) upon PINK1/Parkin-dependent ubiquitination of OMM proteins (Heo et al. 2015; Lazarou et al. 2015; Pickles, Vigié, and Youle 2018). Two such mammalian receptors, Optineurin (OPTN) and p62/Sequestosome-1 (p62/SQSTM1), contain Ubiquitin Binding Domains (UBDs) that facilitate their translocation to the OMM (Kim, Kwon, and Song 2016). While OPTN more uniformly coats the surface of a ubiquitinated mitochondrion (Wong and Holzbaur 2014), p62 is enriched in regions between fragmented mitochondria. Interaction of p62 with poly-ubiquitin is thought to drive phase separation of p62 oligomers and promote clustering of the damaged organelles (Sun et al. 2018). Recruitment and stabilization of OPTN, p62, and other mitophagy receptors on the ubiquitinated organelle promote formation of the isolation membrane by the core autophagy machinery known as ATGs (Rogov et al. 2014; Valverde et al. 2019). Feed-forward mechanisms among mitophagy receptors, ATGs, and the growing double-membrane autophagosome ensure rapid engulfment and sequestration of dysfunctional mitochondria (Chang et al. 2022; Turco, Fracchiolla, and Martens 2020; Vargas et al. 2019), though lysosomal fusion and cargo degradation may occur over longer timescales under physiological conditions (Cason et al. 2021; Evans and Holzbaur 2020).

Mutations in genes encoding key components of the mitophagy pathway are associated with fatal neurodegenerative diseases including Parkinson’s Disease, amyotrophic lateral sclerosis (ALS), and frontotemporal dementia (Ge, Dawson, and Dawson 2020; Menzies et al. 2017; van Rheenen et al. 2021), in keeping with the idea that impaired flux through clearance pathways is deleterious for cellular health. Another major hallmark of these degenerative pathologies is the presence of increasing immune activation over the course of the disease, evidenced by upregulation of innate immune pathways such as the Nuclear Factor kappa B (NF-κB) pathway (Amor et al. 2014; Swarup et al. 2011) and increased levels of cytokines in patient spinal sera (Hu et al. 2017; Nagatsu et al. 2000).

Links between damaged mitochondria and inflammation have already been noted. Cytosolic mitochondrial DNA (mtDNA) is a damage-associated molecular pattern (DAMP) that activates cyclic GMP-AMP synthase-stimulator of interferon genes (cGAS-STING) (West et al. 2015; Yu et al. 2020). Free mtDNA circulating in mitophagy-deficient *PINK1-/-;PRKN-/-* mice sera may directly upregulate production of inflammatory cytokines, initiating or exacerbating degeneration (Sliter et al. 2018). It is also known that dysregulated mitochondrial production of reactive oxygen species (ROS) contributes to activation of the NLR family pyrin domain-containing 3 (NLRP3), which triggers the maturation of pro-inflammatory cytokines (Sarkar et al. 2017; Zhou et al. 2011).

The correlation between neuro-inflammation and exacerbated degeneration supports the idea that abrogating immune responses in patients may be a promising therapeutic strategy. However, small molecule inhibitors of inflammatory pathways have been unsuccessful in mitigating neurodegeneration. More investigation is necessary to identify the roots of deleterious inflammatory activation (Fournier et al. 2018; Wang, Liu, and Zhou 2015).

Here, we identify a novel mechanism for activation of the classical NF-κB pathway by mitophagy intermediates via a Parkin- and p62-dependent mechanism. Following mitophagy initiation and Parkin-dependent ubiquitination of proteins in the mitochondrial outer membrane (OMM), NF-κB Effector Molecule (NEMO) localizes to a subset of depolarized mitochondria with kinetics similar to the recruitment of the well-studied mitophagy receptor OPTN. Rather than promoting mitochondrial clearance, NEMO recruitment triggers multimerization of the Inhibitor of kappa B kinase (IKK) complex, inducing auto-activation that results in the upregulation of NF-κB target genes. Strikingly, mitochondria that recruit NEMO lack the indicators of downstream clearance mechanisms: NEMO-positive mitochondria are less likely to exhibit phospho-OPTN positivity and less likely to be engulfed by ATG8-positive membranes, suggesting that mitophagy and NF-κB signaling may be competing pathways in the response to cellular stress induced by mitochondrial dysfunction.

The induction of NF-κB signaling in response to mitochondrial damage described here parallels some of the cellular responses to bacterial infection (Noad et al. 2017; Van Wijk et al. 2017), suggesting that activation of Parkin-dependent mitophagy has a similar capacity to activate cellular immune responses. We propose that either genetic or environmental factors that induce mitochondrial damage or impair efficient mitophagy may lead to the proliferation of NEMO-positive mitochondria, and thus the activation of IKK signaling and production of disease-associated cytokines, part of the signature neuro-inflammation that can be seen in patients with PD and ALS.

## Results

### NEMO is recruited to damaged mitochondria in a Parkin-dependent manner

OPTN and NEMO each contain an alpha-helical ubiquitin binding domain (UBD) termed the Ubiquitin Binding domain in ABIN and NEMO (UBAN) domain and share 64% sequence homology within their UBAN domains. (Figure 1A) (Li et al. 2016; Yoshikawa et al. 2009). OPTN’s UBAN is required for the efficient recruitment of OPTN to damaged mitochondria following PINK1-dependent activation of the E3-ubiquitin ligase Parkin, which induces the ubiquitination of proteins bound to the OMM (Heo et al. 2015; Wong and Holzbaur 2014). The UBAN of NEMO was initially characterized as a motif required for the recruitment of NEMO to poly-ubiquitination of the activated receptor for the cytokine Tumor Necrosis Factor Alpha (TNFɑ), TNFR1 (Rahighi et al. 2009; Wu et al. 2006). More recent work has shown that NEMO can be recruited to ubiquitinated, cytosolic microorganisms, which act as a signaling platform (Noad et al. 2017; Van Wijk et al. 2017), and that the interactions of NEMO with poly-ubiquitin induce phase separation and activation of the catalytic IKK complex, permitting downstream NF-κB activation (Du et al. 2022).

**Figure 1.**
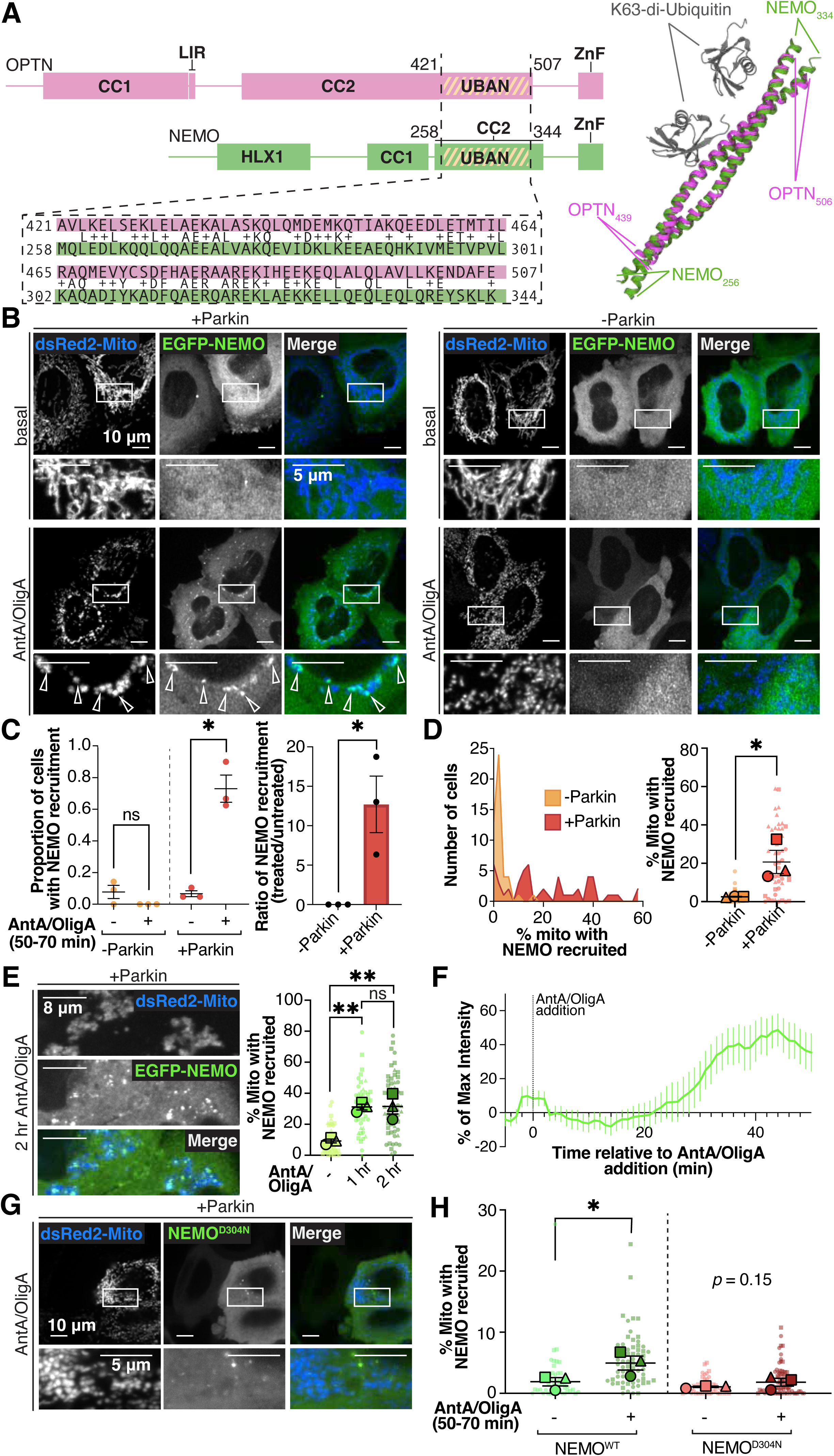
NEMO is structurally related to the mitophagy receptor OPTN and is recruited to damaged mitochondria in a Parkin-dependent manner. (A) Schematics of OPTN (pink) and NEMO (green) proteins with functional and structural domains identified. The UBAN domains of each are highlighted with gold stripes. Below domain maps, the amino acid sequences of the UBANs of OPTN (pink) and NEMO (green) are aligned based on the NCBI BLAST alignment tool. Small lettering between OPTN and NEMO sequences indicates exact residue matches. “+” indicates residues with chemical similarity. (Right) PDB pairwise structure alignment using jFATCAT (flexible) alignment is shown for the UBAN domains of dimerized OPTN (PDB:5WQ4) (pink) and NEMO (PDB: 3JSV) (green), and their interactions with K63-linked di-Ubiquitin (gray). CC1/2, coiled coil domains; LIR, LC3 interacting region; ZnF, zinc-finger domain; HLX1, helical domain. (B) Image of live HeLa cells expressing EGFP-NEMO and a mitochondria-targeted construct (dsRed2-Mito), with (left) or without (right) untagged Parkin. Images were taken at basal conditions (top) or after 50-70 min treatment with 10 μM Antimycin A and 5 μM Oligomycin A (AntA/OligA). Arrows indicate fragmented mitochondria with NEMO recruited. Images are deconvolved and max-projected 2 μm. (C) (Left) Fractions of cells with NEMO recruitment in basal or AntA/OligA conditions. (Right) Ratio of (left) plotted data for AntA/OligA-treated versus basal conditions. Data collected from three independent experiments and analyzed using paired t-tests (left) or Welch’s t-test (right). Error bars indicate standard error of the mean (SEM); ns, not significant; \**P* ≤ 0.05. (D) (Left) Histogram of percent of NEMO-positive mitochondria in a cell, by number of cells observed, and (right) plot of average percentages of NEMO-positive mitochondria. Data collected over 3 separate experiments and analyzed using Welch’s t-test. Error bars indicate SEM; \**P* ≤ 0.05. (E) Images of HeLa cells expressing EGFP-NEMO along with dsRed2-Mito and untagged Parkin. Cells were fixed after 2 hr AntA/OligA treatment. Images are max-projected 2 μm. (Right) Quantification of NEMO occupancy on mitochondria in vehicle conditions, 1 hr, or 2 hr after AntA/OligA addition. Data collected from three independent experiments and analyzed using one way ANOVA with multiple comparisons. Error bars indicate SEM; ns, not significant; \*\**P* ≤ 0.01. (F) Trace of average fluorescent intensity of NEMO recruited to mitochondria over time in 38 events of recruitment after mitochondrial depolarization. Data collected from three independent experiments. Error bars indicate SEM. (See Supp Movie 1). (G) Images of live HeLa cells expressing EGFP-NEMO^D304N^ along with dsRed2-Mito and untagged Parkin after 50-70 min AntA/OligA treatment. Images are max-projected 2 μm. (H) Average percentages of mitochondria per cell with NEMO recruitment. Data collected over 3 separate experiments and analyzed using Welch’s t-test. Error bars indicate SEM; \**P* ≤ 0.05.

We hypothesized that given the homology between NEMO and OPTN, NEMO would be similarly recruited to the ubiquitinated OMM of damaged mitochondria. To test this, we expressed an EGFP-tagged construct of mouse NEMO (EGFP-NEMO) (Tarantino et al. 2014), along with Parkin in HeLa cells (Figure 1B, *left*) and treated the cells with a cocktail of Antimycin A and Oligomycin A (AntA/OligA) in order to depolarize mitochondria (Ordureau et al. 2018). Within 1 hr of AntA/OligA treatment, we observed NEMO puncta resident on depolarized, fragmented mitochondria (Figure 1B, *left, bottom*). This phenomenon was dependent on expression of exogenous Parkin, as HeLa cells do not express endogenous Parkin, and we did not see NEMO recruitment to mitochondria in HeLa cells not transfected with exogenous Parkin (Figure 1B, *right*). We compared the fraction of cells with NEMO puncta after 1 hr AntA/OligA treatment to the fraction with NEMO puncta under basal conditions and found a 13 ± 3.5-fold increase in cells harboring NEMO puncta after mitochondrial damage in the presence of Parkin (Figure 1C). In Parkin-expressing cells, 21 ± 6.0% of mitochondria recruited NEMO 1 hr after AntA/OligA-induced depolarization, while in the absence of Parkin, only 2.5 ± 0.2% of mitochondria were NEMO-positive after AntA/OligA addition (Figure 1D). We observed a range of responses across cells, with up to 60% of mitochondria positive for NEMO in some cells and a less robust response in others. Expression of EGFP-NEMO allowed us to examine the dynamics of recruitment in live cells, but we also noted the recruitment of endogenous NEMO to damaged mitochondria in Parkin expressing cells with immunolabeling (Supp Figure 1A). It is possible that paraformaldehyde (PFA) fixation may enhance the observed number of NEMO puncta in Parkin-expressing, AntA/OligA-treated cells, due to an enhancing effect of PFA on phase separated particles, leading to higher levels than seen in live cell assays (Figure 1E, *1 hr time point*), but in both live and fixed cell assays the recruitment of NEMO to mitochondria was only observed following mitochondrial depolarization.

Next, we tracked individual mitophagy events in Parkin-expressing cells with live cell spinning disk confocal microscopy in order to determine the kinetics of NEMO recruitment (Supp Movie 1). In most of these events, mitochondria gradually accrued NEMO puncta over a period of ∼20 min. However, in ∼25% of events, a preexisting NEMO particle, likely a phase condensate, was recruited to the OMM, leading to a rapid, stepwise increase in fluorescence. We note that cytosolic NEMO particles are present at basal conditions in some cells (Figure 1B, *left cell in basal images*) though they do not appear to associate with the mitochondrial network before depolarization; however, *de novo* NEMO particle formation accounts for the majority of NEMO-positive mitochondria induced by 1 hr AntA/OligA treatment. Occasionally in AntA/OligA conditions, NEMO particles detach from the site of their initial formation and re-attach to a different mitochondrion, suggesting that they maintain their phase-separated nature.

For mitochondria that recruited fluorescently tagged NEMO, the average time to half-maximal fluorescence intensity (half-max) was 37 ± 5.2 min post-AntA/OligA addition (Figure 1F). This result was consistent with our hypothesis that NEMO is recruited by the Parkin-dependent conjugation of poly-ubiquitin, since previous studies showed that Parkin is recruited to mitochondria within 30 min and immediately catalyzes ubiquitin chain linkages at the OMM (Moore and Holzbaur 2016). Damaged mitochondria retained NEMO puncta for at least 5 hours after recruitment, as demonstrated in live and fixed cells (Figure 1E, Supp Figure 1B).

To confirm the role of ubiquitination in NEMO recruitment, we generated a construct in which a conserved residue in the UBAN domain was mutated from aspartic acid to asparagine (NEMO^D304N^), disrupting the ability of NEMO to establish electrostatic interactions with poly-ubiquitin (Rahighi et al. 2009; Wu et al. 2006). Recruitment of NEMO^D304N^ to damaged mitochondria after AntA/OligA treatment was not significantly different than recruitment measured in untreated cells (Figure 1G,H), indicating that the UBAN is required for NEMO recruitment. The limited number of NEMO recruitment events still observed in NEMO^D304N^-expressing cells (Figure 1G,H) may be attributed to homodimerization of NEMO^D304N^ with endogenous, wild-type NEMO. Alternatively, the ZnF domain of NEMO (Figure 1A) may mediate poly-ubiquitin recognition to some extent (Laplantine et al. 2009). However the significantly decreased recruitment of NEMO^D304N^ as compared to NEMO^WT^ in AntA/OligA conditions indicates that the ZnF domain is not sufficient to drive NEMO association with poly-ubiquitinated mitochondria to the same degree. These results demonstrate that NEMO is a novel interactor in the mitochondrial clearance pathway, and its recruitment is dependent on an interaction with ubiquitin.

### NEMO’s interaction with mitochondria is distinct from OPTN

Given the homology in the UBDs of OPTN and NEMO (Figure 1A), we compared their respective occupancies on depolarized mitochondria in Parkin-expressing cells (Figure 2A,B). We found that 18 ± 3.5% of mitochondria recruited only OPTN 1 hr after AntA/OligA addition, while 5.7 ± 1.2% recruited only NEMO, and 6.7 ± 1.5% recruited both OPTN and NEMO (Figure 2B). The remaining ∼70% of mitochondria recruited neither OPTN nor NEMO at this time point; this level of OPTN recruitment is consistent with previous studies (Moore and Holzbaur 2016). These observations demonstrate that OPTN recruitment is more extensive than NEMO, and that mitochondria may recruit either or both of these UBAN-containing proteins.

**Figure 2.**
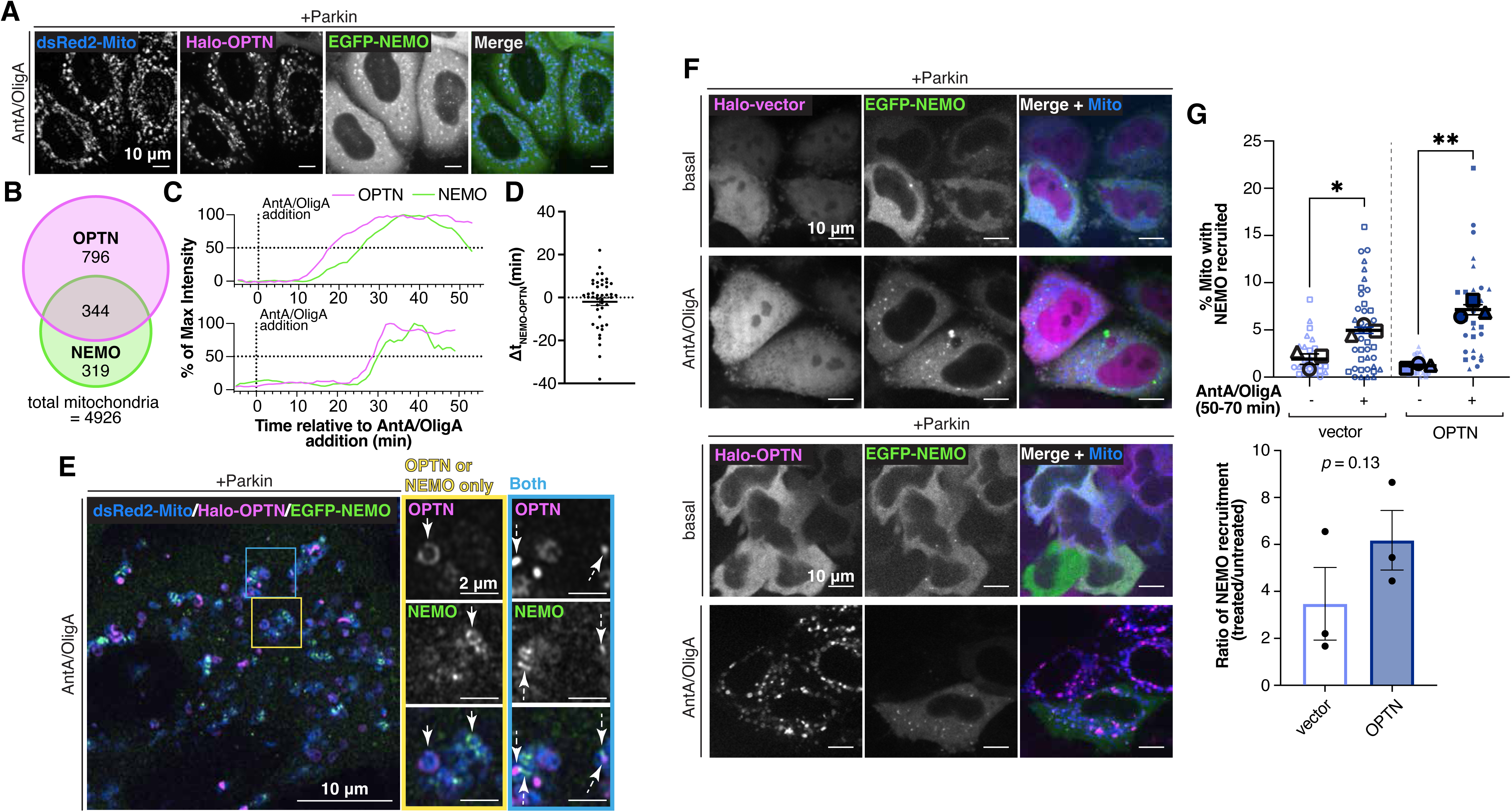
NEMO recruitment is distinct from and independent of OPTN recruitment. (A) Images of live HeLa cells expressing EGFP-NEMO, Halo-OPTN, dsRed2-Mito, and untagged Parkin, treated with AntA/OligA for 50-70 min. Images are max- projected 2 μm. (B) Euler diagram demonstrating the raw number of mitochondria positive for OPTN, NEMO, or both. The remaining mitochondria were positive for neither. (C) Example traces of the relative fluorescent intensities of OPTN and NEMO over time in two events in which both OPTN and NEMO were recruited to a mitochondrion after AntA/OligA-induced depolarization. (See Supp Movie 2) (D) Individual recruitment events were plotted by the difference (minutes) between NEMO and OPTN intensities reaching their respective half-maximums (Δ*t*NEMO-OPTN). Points with negative values indicate events in which NEMO half-max was before OPTN half-max, and vice versa for positive values. Data collected from 16 cells in three independent experiments. Error bars indicate SEM. (E) Airyscan image of a fixed HeLa cell expressing EGFP-NEMO, Halo-OPTN, dsRed2-Mito, and untagged Parkin, treated with AntA/OligA for 60 min. The inset in the yellow box demonstrates examples in which only OPTN (left arrow) or only NEMO (right arrow) were recruited to mitochondria. The blue box demonstrates two examples in which both OPTN and NEMO were recruited to mitochondria (dotted line arrows). (F) Images of live *OPTN-/-* cells expressing EGFP-NEMO, dsRed2-Mito, untagged Parkin, and either Halo-vector (top two rows) or Halo-OPTN (bottom two rows). Cells were imaged in basal conditions (first and third rows) or after 50-70 min AntA/OligA treatment (second and fourth rows). Images are max-projected 2 μm. (G) (Top) Quantification of average percent of mitochondria per cell with NEMO recruitment in cells expressing Halo-vector or Halo-OPTN, in basal or AntA/OligA conditions. (Bottom) Ratios of (top) plotted data, for treated versus control cells. Data were collected over three independent experiments and analyzed using (top) paired t-tests or (bottom) Welch’s t-test. Error bars indicate SEM; \**P* ≤ 0.05; \*\**P* ≤ 0.01.

We then compared the relative timing of OPTN and NEMO recruitment in live cells expressing fluorescently tagged NEMO and OPTN along with a mitochondrial marker, analyzing 43 events in which both NEMO and OPTN were recruited to the same mitochondrion (Supp Movie 2). Neither UBAN-containing protein was consistently recruited first, with instances of NEMO or OPTN reaching half-max from 0 to 40 min before the other. Figure 2C demonstrates examples of two events. On average, NEMO recruitment reached half-max 2.0 ± 1.8 min before OPTN (Δ*t*_NEMO-OPTN_) (Figure 2D). While OPTN intensity increased gradually, in ∼25% of recruitment events we noted a rapid increase in NEMO intensity on depolarized mitochondria, again suggesting NEMO may be recruited as a phase-separated condensate. It is likely that poly-ubiquitin on the OMM recruits both NEMO and OPTN, thus they are recruited in broadly similar time-courses as ubiquitin chains amass on a mitochondrion. However, since OPTN and NEMO are not associated with each other in the cytosol (Zhu et al. 2007), their recruitment kinetics are not tightly correlated.

Next we used Airyscan microscopy to assess the spatial relationship between OPTN and NEMO on individual mitochondria post-recruitment in fixed cells (Figure 2E). We observed that some mitochondria acquired only OPTN or only NEMO (Figure 2E, *yellow box*). However, as described above, some mitochondria demonstrated recruitment of both OPTN and NEMO (Figure 2E, *blue box*). Notably, however, OPTN and NEMO did not overlap on these organelles, but rather occupied discrete, often adjacent domains on a fragmented mitochondrion. Previous work utilizing super-resolution microscopy demonstrated the presence of poly-ubiquitinated microdomains on the surface of cytosolic *Salmonella* (Noad et al. 2017). Fragmented mitochondria may also present these microdomains based on Parkin localization and stabilization on the OMM.

We asked whether OPTN and NEMO compete for poly-ubiquitin on the OMM by testing whether NEMO would be more robustly recruited in the absence of OPTN. We co-expressed Parkin and NEMO in HeLa cells that had been CRISPR-edited to knock out OPTN (*OPTN-/-*) (Lazarou et al. 2015), and rescued these cells with either exogenous OPTN or a Halo-tagged vector (Figure 2F). Upon treatment with AntA/OligA, NEMO was recruited to damaged mitochondria to a similar extent in cells with or without OPTN (Figure 2G, *top*). The fraction of NEMO-positive mitochondria induced by AntA/OligA was not significantly different between vector-expressing and OPTN-expressing cells (Figure 2G, *bottom*). Together, these results indicate that the absence of OPTN does not permit a more extensive recruitment of NEMO to depolarized mitochondria, nor is OPTN is required for NEMO recruitment.

Escape of mitochondrial DNA (mtDNA) to the cytosol has been shown to activate the cGAS/STING pathway of interferon production (West et al. 2015; Yu et al. 2020). Neither NEMO nor the NF-κB pathway has been connected to this response, however we wanted to know whether sites of mtDNA escape would correlate with NEMO puncta. We expressed a fluorescently tagged construct of Transcription Factor A, Mitochondrial (TFAM), which packages mtDNA in the mitochondrial matrix. Almost all TFAM puncta localized to mitochondria, however TFAM location was not a predictor for the location of NEMO puncta on damaged mitochondria (Supp Figure 2B). To account for the possibility that mtDNA comes unbound from TFAM in stressed conditions, we used an antibody to double-stranded DNA (dsDNA), which labels genomic DNA and mtDNA. DsDNA was observed in cell nuclei and in mitochondria, but again, the presence of dsDNA did not correlate with NEMO puncta location (Supp Figure 2C). Together, these results indicated that NEMO recruitment and its patterning on the OMM was regulated by another component.

### p62 is required for NEMO recruitment

The patterning of NEMO recruitment to fragmented, ubiquitinated mitochondria is reminiscent of p62 recruitment: NEMO tends to adopt a bar-like formation between rounded mitochondria (Figure 2E, *blue box*) much like p62 occupancy at the interstices of damaged mitochondria (Sun et al. 2018; Wong and Holzbaur 2014). We co-expressed Parkin, NEMO, and p62 in HeLa cells CRISPR-edited to knock out p62 (*p62-/-*) (Sarraf et al. 2020) and found that upon mitochondrial depolarization, NEMO and p62 demonstrated striking colocalization to fragmented mitochondria (Figure 3A,B). We found the same coincidence of NEMO and p62 when we probed for endogenous p62 in wild-type HeLa cells (Supp Figure 3A). We conducted live imaging on cells co-expressing Parkin, p62, and NEMO along with a mitochondrial marker and compared the recruitment kinetics of p62 and NEMO in 16 separate events (Supp Movie 3, Figure 3C). Unlike the wide distribution of Δ*t* between NEMO and OPTN recruitment (Figure 2D), NEMO and p62 were recruited with very similar kinetics, with a smaller average Δ*t*_NEMO-p62_ and a smaller variance among events (-1.6 ± 1.1 min) (Figure 3D). Together, these observations indicate that the recruitment of NEMO and p62 to depolarized mitochondria are correlated in both space and time.

**Figure 3.**
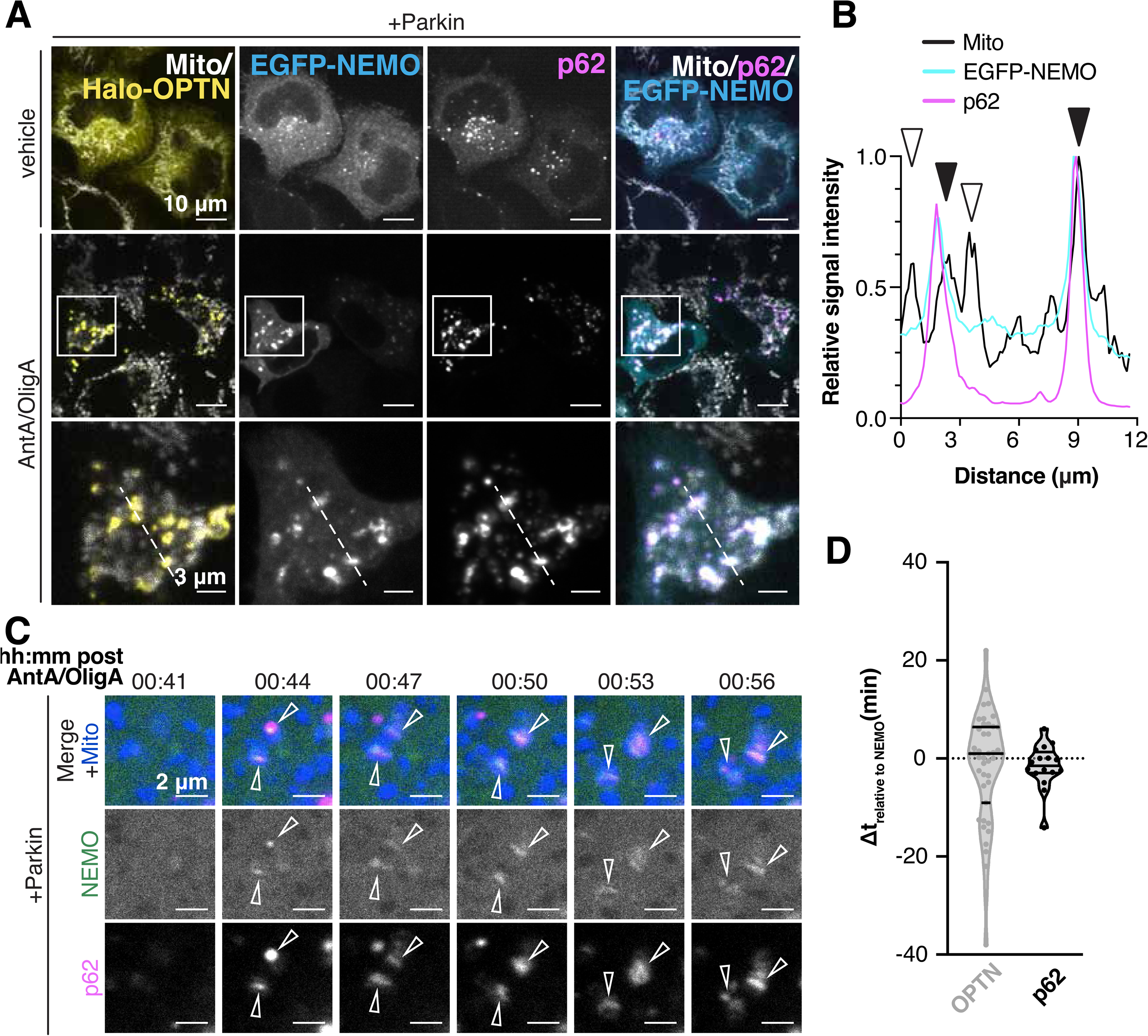
NEMO and p62 recruitment to depolarized mitochondria are spatially and temporally linked. (A) Images of *p62-/-* cells expressing EGFP-NEMO, Halo-OPTN, mCherry-p62, and untagged Parkin, fixed after treatment in the respective conditions and labeled with antibodies to p62 and HSP60 (Mito). Dotted white line indicates where line scan data was collected (B). Images are max-projected 2 μm. (B) Line scan of relative fluorescence intensities of Mito, NEMO, and p62 in the AntA/OligA-treated cells in (A). Solid arrows indicate mitochondria with recruitment of colocalized NEMO and p62. Unfilled arrows indicates mitochondria with no recruitment of NEMO or p62. (C) Stills taken from live, confocal time-lapse images of HeLa cells expressing EGFP-NEMO, mCherry-p62, a mitochondria-targeted BFP2 construct (Mito), and untagged Parkin over the course of AntA/OligA treatment. Time post-AntA/OligA is indicated above each still. Arrows mark two events in which p62 and NEMO were recruited to mitochondria. (See Supp. Movie 3) (D) Individual recruitment events were plotted by the difference (minutes) between NEMO and OPTN intensities (data copied from 2D) or NEMO and p62 intensities reaching their respective half-maximums (Δ*t*_NEMO-OPTN_). Points with negative values indicate events in which NEMO half-max was before OPTN or p62 half-max, and vice versa for positive values. Data for p62 recruitment were collected from six cells from three independent experiments. Solid lines within violin plots indicate median and first and third quartiles.

We then asked whether p62 is required for NEMO recruitment to depolarized mitochondria. Depletion of p62 from wild-type HeLa cells using siRNA resulted in a significant diminishment of NEMO recruitment upon AntA/OligA application compared to mock siRNA-treated cells in the same condition (Figure 4A-C), indicating that p62 is required for NEMO recruitment to damaged mitochondria. There are several well-characterized domains in p62 (Figure 4D), including an N-terminal (1-102) PB1 domain that promotes multimerization with other PB1-domain containing proteins and a C-terminal alpha-helical UBA-type UBD (389-434) that mediates interactions with poly-ubiquitin chains. Much of the middle segment of the protein is a disordered region with motifs that promote interactions with a number of signaling molecules, including TNF Receptor Associated Factor 6 (TRAF6) (TIR) and LC3 (LIR) (Jakobi et al. 2020). We expressed a series of mutated and truncated p62 constructs (Linares et al. 2013; Wurzer et al. 2015) (Figure 4D) in *p62-/-* cells and found that NEMO recruitment to damaged mitochondria mirrored p62 recruitment in each case (Figure 4D). Mutations to the PB1 domain of p62 abrogate the ability of p62 to oligomerize and form filaments, to the detriment of its ability to be recruited to poly-ubiquitin (Wurzer et al. 2015). Thus, a deletion of the PB1 domain (ΔPB1) or two point mutations of lysine to alanine and aspartic acid to alanine within the PB1 domain (PB1^AA^) resulted in significantly fewer mitochondria positive for NEMO as compared to wild-type p62 expression (Figure 4D,E). Expression of a p62 mutation with a deletion of the UBA (ΔUBA) also precluded p62 recruitment to damaged mitochondria and suppressed the fraction of NEMO-positive mitochondria. In contrast, mutations to p62 that did not affect its recruitment to damaged mitochondria (LIR^AAAA^ and TIR^AAA^) did not significantly alter NEMO recruitment (Figure 4D-E). The recruitment of OPTN was not affected by expression of any of the p62 variants (Figure 4E, *left column*)

**Figure 4.**
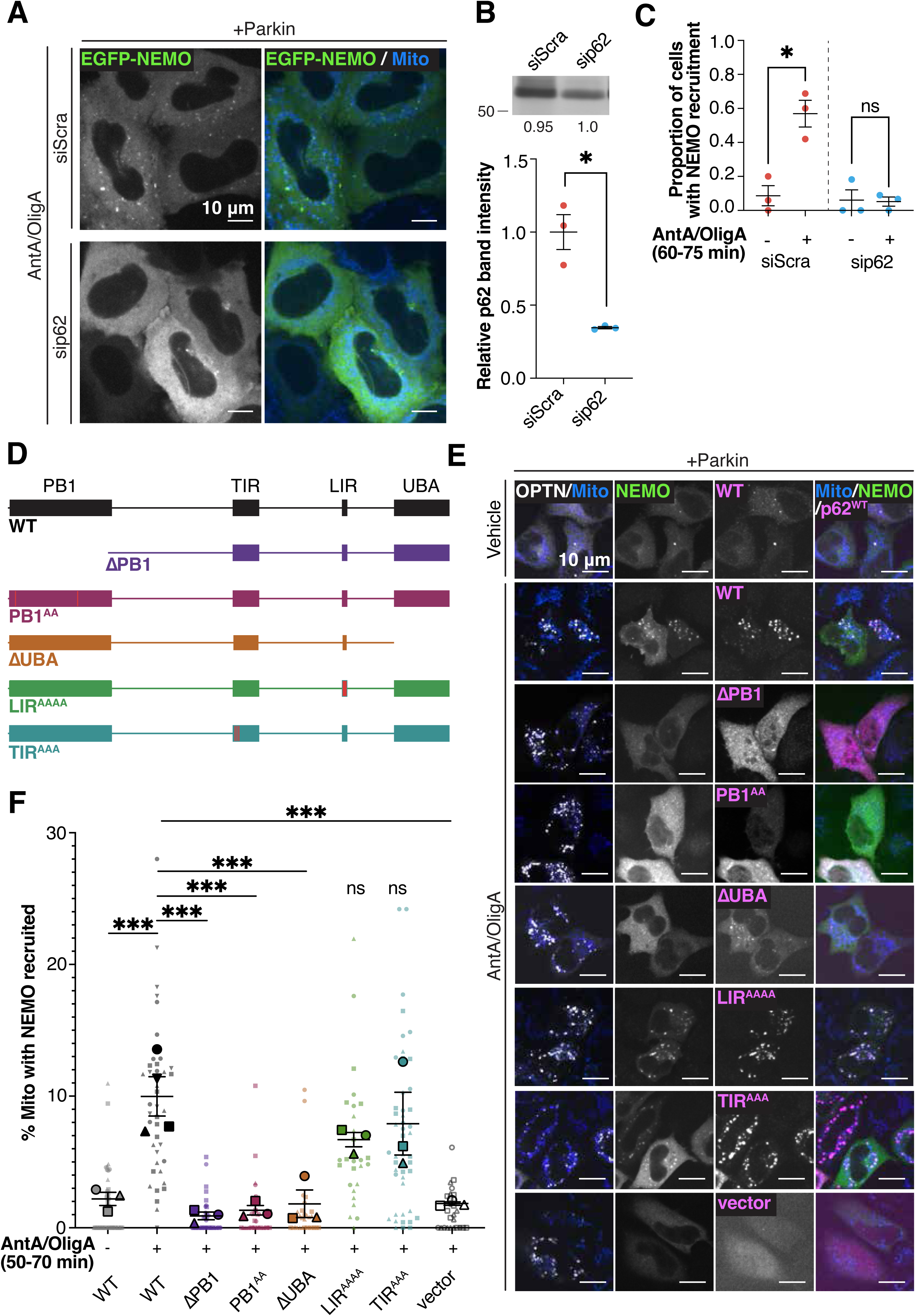
NEMO recruitment to depolarized mitochondria is dependent on recruitment of oligomerized p62 (A) Images of live HeLa cells expressing EGFP-NEMO, dsRed2-Mito and untagged Parkin, treated with AntA/OligA for 50-70 min. Cells expressed a scrambled control siRNA (top row) or p62-targeted siRNA (bottom row). Images are max-projected 2 μm. (B) Representative Western blot for p62 from HeLa cells expressing a scrambled control siRNA or p62-targeted siRNA. Loading control amounts indicated below each lane. Plot shows the p62 band intensity relative to total protein for three experiments. Data were collected over three independent experiments and analyzed using paired t-test. Error bars indicate SEM; \**P* ≤ 0.05. (C) Quantification of cells categorized by whether there was NEMO recruitment or not in vehicle or AntA/OligA conditions. Data were collected over three independent experiments and analyzed using paired t-tests. Error bars indicate SEM; ns, not significant; \**P* ≤ 0.05 (D) Domain maps for p62 variants expressed in *p62-/-* cells. Red bars indicate locations of alanine substitutions. PB1, Phox and Bem1 domain; TIR, TRAF6 interacting region; LIR, LC3 interacting region; UBA, Ubiquitin-assotiated domain. (E) Images of *p62-/-* cells expressing EGFP-NEMO along with, Halo-OPTN, untagged Parkin, and one of six mCherry-p62 constructs or the mCherry vector control, fixed after treatment in the respective conditions and labeled with antibodies to p62 and HSP60 (Mito). Images are max-projected 2 μm. (F) Quantification of the extent of NEMO recruitment to mitochondria in *p62-/-* cells expressing the respective constructs. Data were collected over three independent experiments and analyzed using one way ANOVA with multiple comparisons. Error bars indicate SEM; ns, not significant; \*\*\**P* ≤ 0.001.

To determine whether NEMO and p62 are associated in the cytosol and thus recruited as a preexisting co-complex, we tested whether soluble NEMO would co-precipitate with p62 (Supp Figure 4A). p62 demonstrated a high degree of non-specific binding in this assay, however there was no evidence to support an interaction between NEMO and p62 in cytosolic extracts from cells treated with AntA/OligA or a vehicle control. Next, we expressed EGFP-tagged ubiquitin and tested the co-precipitation of p62 and NEMO from *p62-/-* cells as performed previously (Supp Figure 4B) (Wurzer et al. 2015). As expected, addition of wild-type p62 resulted in p62 co-precipitation with EGFP-Ubiquitin, while expression of p62^ΔPB1^ resulted in much less p62 associated with ubiquitin. Neither expression of wild-type p62 nor p62^ΔPB1^ resulted in co-precipitation with NEMO. Together these results indicate that recruitment and oligomerization of p62 are necessary but not sufficient for NEMO recruitment.

### NEMO recruitment occurs separately from the autophagosome expansion pathway

Since OPTN, p62, and NEMO demonstrate distinct patterns of recruitment and occupancy on the OMM after mitochondrial depolarization, we tested whether there were downstream consequences depending on which UBD-containing proteins were recruited. OPTN phosphorylation is catalyzed by TANK-binding kinase 1 (TBK1) and Unc-like Autophagy Activating Kinase 1 (ULK1), and is required for efficient clearance of damaged mitochondria (Harding et al. 2021; Moore and Holzbaur 2016; Richter et al. 2016). We probed phospho-OPTN with an antibody specific for the phosphorylation of OPTN residue serine-177 (pOPTN). As expected, Parkin-expressing cells that had undergone depolarization damage exhibited many pOPTN-positive mitochondria (Figure 5A). Notably, these mitochondria tended to be distinct from the population of mitochondria that recruited NEMO (Figure 5A,B), with only a small fraction of mitochondria recruiting both.

**Figure 5.**
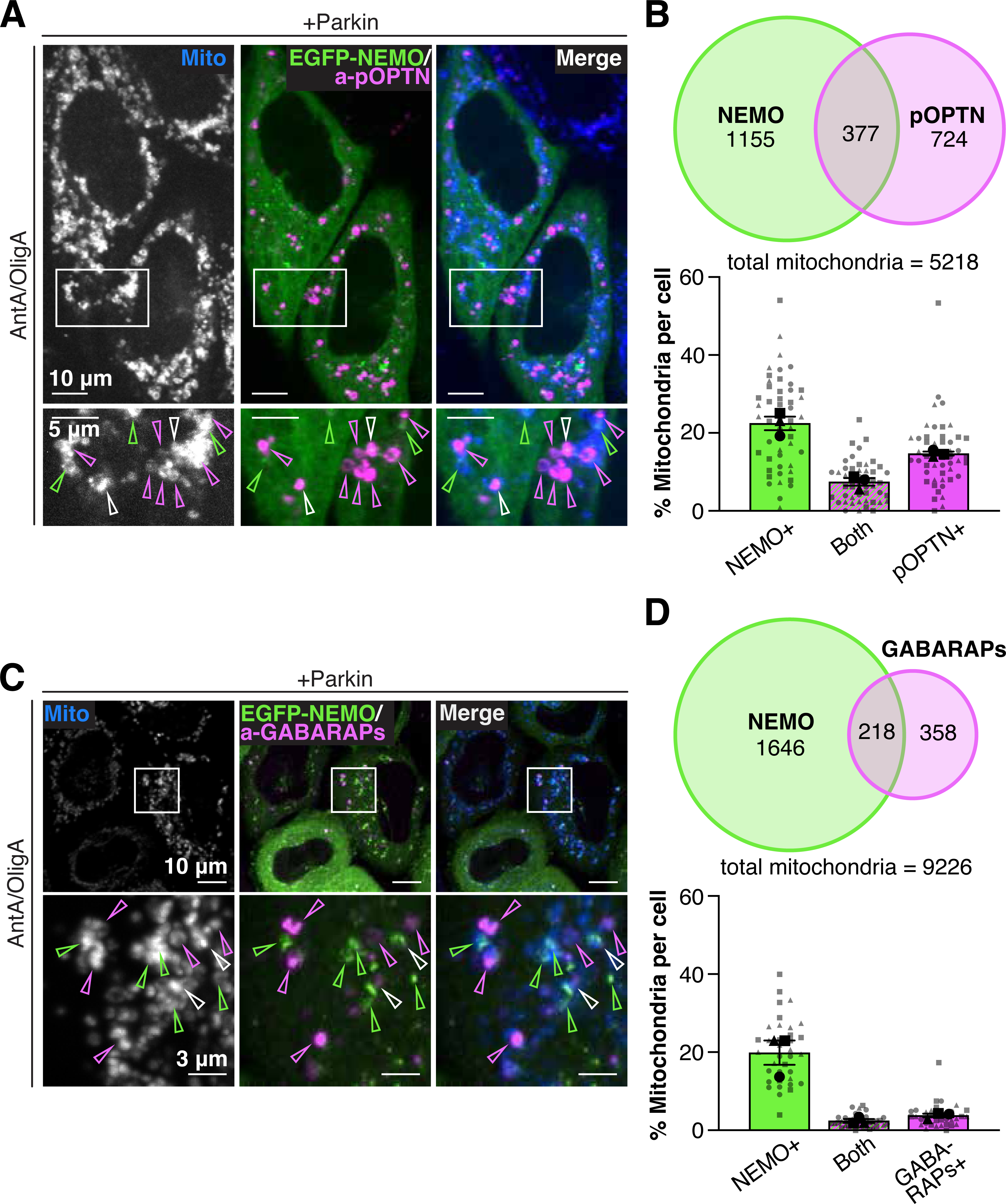
NEMO and LC3/GABARAPs are recruited to distinct mitochondrial populations. (A) Images of HeLa cells expressing EGFP-NEMO along with dsRed2-Mito and untagged Parkin, fixed after AntA/OligA treatment and labeled with an antibody to phospho-S177-OPTN (pOPTN). In zoom images (bottom row), arrows indicate mitochondria with only NEMO recruitment (green) only pOPTN (magenta) or both (white). Images are max-projected 2 μm. (B) (Top) Euler diagram demonstrating the raw number of mitochondria positive for NEMO, pOPTN, or both. The remaining mitochondria were positive for neither. (Bottom) Average percent of mitochondria per cell positive for only NEMO, only pOPTN, or both. Data collected from 3 independent experiments. (C) Images of HeLa cells expressing EGFP-NEMO along with dsRed2-Mito and untagged Parkin, fixed after AntA/OligA treatment and labeled with an antibody to GABARAP, GABARAPL1, and GABARAPL2 (GABARAPs). In zoom images (bottom row), arrows indicate mitochondria with only NEMO recruitment (green) only GABARAPs (magenta) or both (white). Images are max-projected 2 μm. (D) (Top) Euler diagram demonstrating the raw number of mitochondria positive for NEMO, GABARAPs, or both. The remaining mitochondria were positive for neither. (Bottom) Average percent of mitochondria per cell positive for only NEMO, only GABARAPs, or both. Data collected from 3 independent experiments.

The distinction we noted between subpopulations of mitochondria positive for either NEMO or pOPTN made us wonder whether NEMO recruitment would also be negatively correlated with markers of downstream steps of mitophagy. ATG8 family proteins are required for the formation of the autophagosomal membrane (Tanida, Ueno, and Kominami 2004; Turco, Fracchiolla, and Martens 2020). The family comprises six members of small, ubiquitin-like proteins: LC3A, LC3B, LC3C, GABARAP, GABARAPL1, and GABARAPL2. In both selective and non-selective autophagy, assembly of ATG8-positive membranes indicates engulfment of cellular material for degradation (Tanida, Ueno, and Kominami 2004). We probed Parkin-expressing cells with an antibody for GABARAP, GABARAPL1, and GABARAPL2 (GABARAPs) and found that after mitochondrial depolarization, only a small population (<10%) of fragmented mitochondria were positive for both NEMO and GABARAPs; instead, almost half of mitochondria were positive for either NEMO or the ATG8s while half recruited neither at this time point (Figure 5C,D). This result was consistent with a parallel experiment in which we measured recruitment of NEMO and LC3B (Supp Figure 5A). The partitioning of mitochondria into sub-populations that recruited either NEMO or markers of downstream clearance suggests that mitochondria not engulfed by the autophagosome maintain persistent NEMO localization.

### The activated IKK complex promotes NF-ĸB signaling from damaged mitochondria

NEMO is an essential regulator of classical NF-ĸB signaling; specifically, NEMO is one of three subunits that may comprise the IKK complex. While NEMO is a structural, non-catalytic component, the two other subunits, IKKɑ and IKKβ are kinases, which are activated via phosphorylation of IKKβ upon multi-IKK complexing and phase separation (Du et al. 2022; Ko et al. 2022; Rushe et al. 2008). In order to determine whether IKKβ is co-recruited to damaged mitochondria with NEMO, we co-expressed NEMO with a fluorescently tagged construct of IKKβ in the presence of Parkin (Figure 6A). Upon depolarization of mitochondria by AntA/OligA, there was colocalization of NEMO and IKKβ at fragmented mitochondria, indicating IKK multi-complex formation (Figure 6A, *right*). We also found that endogenous IKKβ was recruited to damaged mitochondria by staining in fixed cells (Supp Figure 6A).

**Figure 6.**
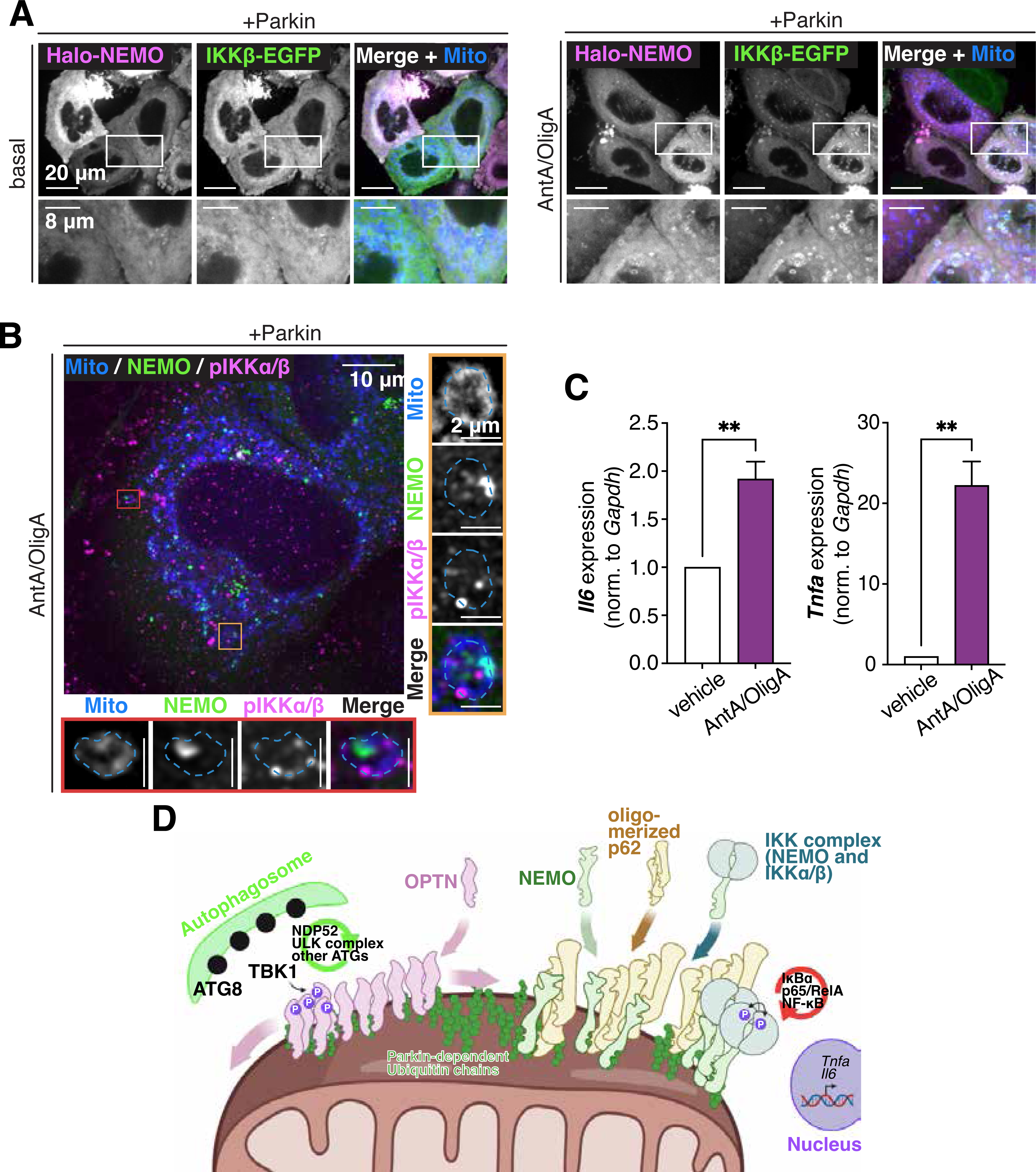
The IKK complex is recruited to damaged mitochondria, which serve as a platform for activation of a NF-κB signaling. (A) Images of live HeLa cells expressing Halo-NEMO, IKKβ-EGFP, dsRed2-Mito and untagged Parkin. Images are deconvolved and max-projected 2 μm. (B) Image of a HeLa cell expressing EGFP-NEMO and untagged Parkin, fixed after AntA/OligA treatment and labeled with antibodies to TOM20 (Mito) and phospho-IKKɑ/β. Zoom images (red and gold boxes) show individual mitochondria with NEMO recruitment and IKK activation. Blue dotted line indicates mitochondrion outline. Image is deconvolved and max-projected 2 μm. (C) RT-qPCR results for HeLa cells expressing Parkin and EGFP-NEMO treated with vehicle or AntA/OligA. Transcripts for *Il6* and *Tnfa* were measured relative to those for *Gapdh*. Data was collected from 3 independent experiments. (D) Model depicting a depolarized mitochondria acting as a signaling platform for the activation of the IKK complex via NEMO recruitment. Simultaneously, OPTN recruitment mediates the expansion of the ATG8-positive double-membrane autophagosome in order to engulf the damaged organelle. We hypothesize NF-κB signaling and mitophagy are competing pathways regulating the response to cellular stress.

NEMO recruitment and binding to poly-ubiquitin chains is thought to induce phase separation of the IKK complex and conformational changes among the IKK subunits, both of which promote IKKβ phosphorylation and complex activation (Du et al. 2022; Ko et al. 2022). We probed fixed cells with an antibody to phospho-Ser176/pSer180-IKKα/β and found that mitochondria positive for NEMO recruitment also demonstrated evidence of activated IKK (Figure 6B). Together these data suggest that NEMO is co-recruited to damaged mitochondria with the catalytic IKK components, and the complex is activated via phosphorylation.

In canonical NF-ĸB signaling, active IKK induces the phosphorylation of IĸBɑ, promoting its degradation and permitting nuclear translocation of NF-ĸB transcription factors. Given the recruitment and activation of the IKK complex at damaged mitochondria, we wanted to know whether NF-ĸB target genes would be upregulated in the context of Parkin-dependent mitophagy. We collected mRNA from Parkin-expressing cells in basal conditions or after AntA/OligA treatment and performed RT-qPCR to detect the levels of transcripts encoding the pro-inflammatory cytokines interleukin 6 (IL6) and TNFɑ. AntA/OligA-treated cells produced significantly higher levels of *Il6-* and *Tnfa-* encoding mRNA (Figure 6C). In contrast, we did not find evidence of induction of an interferon-type response characterized by upregulation of antiviral transcripts *Ifit1* or *Ifit3* (Supp Figure 6B). This pattern is consistent with induction of an innate immune response that promotes microglial activation and proliferation (Chiu et al. 2013), features documented repeatedly in patients with neurodegeneration (Philips and Robberecht 2011; Sitte et al. 2001). The lack of IFIT upregulation corroborates our finding that cells are not undergoing a cGAS-STING response to cytosolic mtDNA in the context of our assays (Supp Figure 2) (West et al. 2015). Together, our data demonstrate a novel mechanism by which depolarized, poly-ubiquitinated mitochondria recruit the NF-ĸB effector NEMO and promote inflammatory signaling (Figure 6D).

## Discussion

Mitochondria have been frequently implicated in innate immunity, ranging from ontology studies linking mitochondrial quality control genes to autoimmune diseases, to the discovery that mtDNA is a Damage-Associated Molecular Pattern (DAMP). Here, we characterize a novel mechanism by which damaged mitochondria in Parkin-expressing cells become intracellular platforms that provoke classical NF-ĸB signaling via NEMO, simultaneous to their recruitment of OPTN and activation of mitophagy (Figure 6D). Our model is reminiscent of *Salmonella* Typhimurium invasion of the cytosol, in which ubiquitin microdomains on the exposed bacterial surface recruit both xenophagic elements and the IKK complex, activating NF-ĸB signaling (Noad et al. 2017; Van Wijk et al. 2017). In the case of cytosolic bacteria, autophagosomal clearance and NF-ĸB activation are independently important to cell health and viability. However, given that mitochondrial dysfunction and neuro-inflammation are hallmarks of neurodegenerative disease, we propose that chronic activation of the mitochondrial signaling paradigm elaborated here is deleterious and contributes to progressive degeneration.

First, we documented Parkin-dependent NEMO recruitment to damaged mitochondria with similar kinetics to the mitophagy receptor OPTN (Figure 1). Recent work has demonstrated that interactions between NEMO and poly-ubiquitin drive phase separation of the IKK complex into condensates as part of NF-ĸB activation (Du et al. 2022). We used Airyscan technology to image NEMO and OPTN on the OMM and found that strikingly, while OPTN is typically organized into an apparently homogenous ring around fragmented mitochondria, NEMO forms irregular puncta on the OMM that rarely envelop the organelle (Figure 2E). The NEMO puncta we observed may be liquid droplets that associate with mitochondria due to the abundance of ubiquitin chains deposited by Parkin on the OMM (Ordureau et al. 2020). When they did coincide on a single organelle, NEMO and OPTN occupied segregated domains on the OMM, suggesting that NEMO bodies exclude OPTN. We found that NEMO-positive domains were not correlated with TFAM or mtDNA, but that NEMO occupancy was strongly correlated with the localization of another mitophagy receptor, p62 (Figures 3, 4). Indeed, p62 recruitment to mitochondria was required for NEMO recruitment (Figure 4). The topography of ubiquitin linkages on the OMM may contribute to the stability of either OPTN or NEMO-p62 residence. Parkin ligase activity generates mainly K63 chains (Ordureau et al. 2018), however, longer or shorter chains may result in preferential recruitment of various receptors. Further work must be done to elucidate the nature of the segregated microdomains on the OMM.

Since p62 also forms condensates in a manner dependent on its PB1 domain and interactions with poly-ubiquitin (Jakobi et al. 2020; Sun et al. 2018), p62 may be incorporated with NEMO into phase separated droplets associated with the OMM. Direct p62-NEMO interactions have been reported in studies of NF-ĸB activation upon TNFɑ and IL-1β stimulation (Schimmack et al. 2017; Zotti et al. 2014), but not in the context of mitochondrial clearance. Previously, Rose et. al. performed tandem mass tag spectrometry to determine ubiquitinated targets in a Dox-inducible Parkin system. As expected, OPTN and TBK1 were among the highly ubiquitinated targets within 1 hr of AntA/OligA treatment. After 6 hr, p62 and NEMO were highly ubiquitinated, while the other receptors were less represented in the sample pool (Rose et al. 2016). This could indicate that a portion of mitochondria not cleared in the first 6 hr after depolarization accumulate NEMO and p62.

Although previous studies have shown that OPTN can out-compete NEMO for poly-ubiquitinated Receptor-Interacting Protein (RIP) at the TNFR1 (Zhu et al. 2007), we found that NEMO recruitment was not amplified by depletion of OPTN (Figure 2F,G). Instead, we propose that there is a “ceiling” for NEMO residence on the OMM of damaged mitochondria, which is potentially correlated to p62 availability or another unidentified factor specific to the mitophagy context. Other mediators of classic NF-ĸB signaling such as TAK1 and TAB1/2 may also be recruited. It may be beneficial for cells to preserve a limit to mitochondrial NEMO accumulation in order to dampen potential downstream signaling in the event of isolated stress events. However, we hypothesize that the chronic localization of NEMO and the active IKK complex is deleterious to cellular homeostasis, permitting prolonged NF-ĸB signaling and upregulation of cytokine expression.

The potential for mitophagy intermediates to evoke NEMO-mediated NF-ĸB signaling has implications for dysfunctional clearance pathways associated with neurodegeneration. Parkin is expressed in neural cell types, such as neurons, microglia, and astrocytes (Evans and Holzbaur 2019; Zhang et al. 2016), and loss of function mutations in both *PINK1* and *PRKN* genes cause adult-onset PD (Ge, Dawson, and Dawson 2020), suggesting that Parkin-mediated mitophagy is a critical homeostatic mechanism for the brain. We previously showed that neurons expressing TBK1 mutants associated with ALS have decreased mitochondrial membrane potential and a greater extent of mitophagy even under basal conditions (Harding et al. 2021). An overburdened clearance pathway could result in an enhancement of NEMO-mediated NF-ĸB signaling over time, leading to upregulation of cytokines, as seen in the RT-qPCR assay reported here and in the cerebrospinal fluid of patients with neurodegenerative disease (Hu et al. 2017).

On the other hand, it is important to note that NF-ĸB signaling is crucial to cell survival pathways, promoting the expression of anti-apoptotic proteins such as calbindin and Bcl-2 family members (Mattson et al. 2000). Parkin-dependent NEMO recruitment and IKK activation may be a built-in brake to neuronal death pathways triggered by mitochondrial failure. In cases of Parkin deficiency, such as PD, Parkin-independent clearance pathways may be sufficient to stave off deleterious effects initially, but over the lifetime of an organism, the absence of mitochondrial poly-ubiquitination and NEMO signaling in neurons could ultimately permit degeneration and cell death. Overall, NEMO-mediated pathways for inflammation and cell survival are likely tightly regulated, whether in the case of invasive bacteria or malfunction of resident bacterial descendants, the mitochondria (Van Wijk et al. 2017; Youle 2019). Any such signaling activated by mitophagy intermediates must be accounted for as we investigate underlying mechanisms of neurodegenerative pathology.

## Materials and Methods

### Reagents

Constructs used were: Mito-DsRed2 (kindly provided by T. Schwarz, Harvard Medical School, Boston) and Mito-SBFP2 (Addgene #187964); untagged Parkin (Addgene #187897); EGFP-NEMO (kindly provided by E. Laplantine, Institut Pasteur, Paris), EGFP-NEMO^D304N^ (Addgene #187898), and Halo-NEMO (Addgene #187899); Halo-OPTN (Addgene #187900); mCherry constructs, vector (Addgene #187983), WT (Addgene #187986), ΔPB1 (Addgene #187985), PB1^AA^ (Addgene #187982), ΔUBA (Addgene #187984), and LIR^AAAA^ (Addgene #187981) were kindly provided by S. Martens, University of Vienna, Austria, and TIR^AAA^ (Addgene #187901) was generated by site-directed mutagenesis of mCherry-p62^WT^); IKKβ (aka IKK2)-EGFP (Addgene #111195); TFAM-SNAPf (Addgene #187909); and EGFP-Ubiquitin (Addgene #187910). siRNA targeting p62, ON-TARGETplus p62 siRNA (J-010230-05, target sequence GAACAGAUGGUCGGAUA) and scrambled control siRNA were from Horizon Discovery. SNAP ligand (SNAP-Cell 647-SiR, S9102S) was from New England BioLabs. Halo ligand (JaneliaFluor 646, GA112A) was from Promega. Antibodies used were: anti-HSP60 (Sigma Aldrich, SAB4501464, 1:50), anti-TOM20 (Santa Cruz, sc-17764, 1:50), anti-p62 (abcam, ab56416, IF: 1:250, WB: 1:500), anti-phospho-OPTN (Ser177) (Cell Signaling Technology, 57548, 1:200), anti-GABARAP+GABARAPL1+GABARAPL2 (abcam, ab109364, 1:125), anti-LC3B (abcam, ab109364, 1:50), anti-phospho-IKKα/β (Ser176/180) (Cell Signaling, #2697, 1:100), anti-IKKβ (Novus, NBP2-33214, 1:100); anti-NEMO (abcam, ab178872, IF: 1:250, WB: 1:5000), anti-dsDNA (Millipore-Sigma, CBL186, 1:50), anti-GFP (abcam, ab290, 1:375), anti-GAPDH (abcam, ab8245, 1:500). The drugs Antimycin A (A8674) and Oligomycin A (75351) were purchased from Sigma-Aldrich. Primers used for RT-qPCR were IFIT1 (forward AGCCTAGAGGGCAGAACAGA, reverse GCCAGGTCTAGATGAGCCAC), IFIT3 (forward TGCAGGGAAACAGCCATCAT, reverse GGCATTTCAGCTGTGGAAGG), IL6 (forward AGGAGCCCAGCTATGAACTC, reverse GAGAAGGCAACTGGACCGA), TNFA (forward GGCCCGACTATCTCGACTTT, reverse CTCACAGGGCAATGATCCCA), and GAPDH (forward ATCTTCTTTTGCGTCGCCAG, reverse GTTGACTCCGACCTTCACCT).

### Cell culture and transfection

Cells used: HeLa-M, a variant of HeLa cells (Caviston et al. 2007), were a generous gift from A. Peden, University of Sheffield, U.K.; HeLa-*OPTN-/-* (Lazarou et al. 2015), and HeLa-*p62-/-* (Sarraf et al. 2020). All cells were maintained in DMEM (Corning, 10-017-CV) with 10% fetal bovine serum (HyClone) and 1% GlutaMAX glucose supplement (Gibco, 35050061). Cells were maintained in an environment of 37°C with 5% CO2. Passage number was below 30 for HeLa-M cells and below 15 for knock-out lineages. Each line was tested for mycoplasma regularly. 18-20 hours prior to fixation, live imaging, or collection, cells were approximately 80-90% confluent, and were transfected with the appropriate constructs using Lipofectamine 2000 (ThermoFisher Scientific, 11668027). For p62 knockdown, cells were transfected with 20 μM sip62 or siCtrl and other constructs when they were 80-90% confluent and imaged 40-48 hr post-transfection.

### Labeling and treatment performed on live cells

To label Halo-tagged proteins, cells were incubated with 190 nM Halo ligand in media for at least 20 min. For SNAP-tagged proteins, cells were incubated with 1.25 μM SNAP ligand in media for at least 45 min. For depolarization (or vehicle control), cells were treated with a combination of 5 μM Oligomycin A (or DMSO) and 10 μM Antimycin A (or EtOH).

### Fixation

Cells were fixed with 4% paraformaldehyde (PFA). For imaging of autophagosomes (GABARAPs and LC3B), after PFA fixation cells were permeabilized in methanol and blocked in 5% goat serum, 1% BSA. For staining pIKK and IKKβ, after PFA fixation cells were permeabilized with 0.2% Triton-X and blocked in 10% FBS in DMEM (Paul and Schaefer 2015). For all others, cells were permeabilized in 0.5% Triton-X and blocked with 3% BSA, 0.2% Triton-X. Cells were incubated with primary and secondary antibodies in appropriate blocking buffers.

### Microscopy

For live cell imaging, conditioned media was replaced with Leibovitz’s L-15 Medium (Gibco, 11415064) supplemented with 10% fetal bovine serum. Fields of view were chosen to maximize the number of healthy-appearing cells that expressed detectable components of interest. All samples except where indicated as AiryScan were imaged with a Nikon Eclipse Ti Microscope with a 60X objective (Apochromat, Nikon, 1.40-N.A. oil immersion) or 100X objective (Apochromat, Nikon, 1.49-N.A. oil immersion) and an UltraView Vox spinning disk confocal system (PerkinElmer) with a CMOS ORCA-Fusion camera (Hamamatsu, C11440-20UP). Z-stacks were collected for all samples except timelapses. The parameters for Z-stacks were 0.15 nm/step through at least 3.1 μm of cells’ midsections. Timelapse imaging was collected for confocal sections at 60 seconds/frame. For timelapse imaging with the addition of AntA/OligA, 5-10 frames were collected at basal conditions, then a volume of imaging media at least 50% of the initial volume was added, including AntA/OligA, to bring the total concentration to 10 μM/5 μM as frame collection continued. Volocity (PerkinElmer) or Visiview (Visitron Systems) were used as acquisition softwares. For AiryScan detection (Figure 2E), fixed samples were imaged with with a Zeiss Axio Observer inverted microscope with a 63X objective (oil immersion). Data were collected with ZEN (Zeiss) v2.3 acquisition software.

### Co-immunoprecipitations and immunoblots

For standard cell lysis (Figure 4B), cells were lysed with RIPA buffer (50 mM Tris-HCl, 1 mM EDTA, 2 mM EGTA, 1% Triton X, 0.5% sodium deoxycholate, 0.1% sodium dodecyl sulfate, 150 mM NaCl) with added Halt Protease and Phosphatase Inhibitor Cocktail (ThermoFisher Scientific, 78444) and assayed for protein concentration with Pierce BCA Protein Assay Kit (ThermoFisher Scientific, 23225). For GFP-NEMO immunoprecipitation, cells were lysed by freeze-thaw and suspended in buffer as described previously (Turco et al. 2019). For GFP-Ubiquitin immunoprecipitation, cells were lysed as described previously (Wurzer et al. 2015). GFP-Trap Magnetic Particles (Chromotek M-270) were used to immunoprecipitate GFP conjugated elements. All samples were assayed by SDS-PAGE and labeled with fluorescent secondary antibodies (Li-Cor) for imaging on an Odyssey CLx machine (Li-Cor).

### RNA extraction and RT-qPCR

∼One million cells were collected for each condition after 5 hr incubation with vehicle control or AntA/OligA using standard TRIzol protocol (ThermoFisher, 15596026) to extract RNA in cells expressing Parkin and EGFP-NEMO. cDNA was generated using the SuperScript First-Strand Synthesis System (ThermoFisher, 11904018). cDNA was cleaned using Zymo Research Oligo Clean & Concentrator kit (D4060). 11 ng samples or equivalent of nuclease free water for No Transcript Control were added to each reaction with 300 nM of each primer. PowerUP SYBR Green Master Mix (Applied Biosystems, A25742) was used to catalyze PCR in a QuantStudio 3 Real-Time PCR System Machine (Applied Biosystems, A28567). Amplification data was produced with QuantStudio Design and Analysis software.

### Image processing and data analysis

Microscopy images were deconvolved where indicated with Huygen’s Professional version 17.10 software (Scientific Volume Imaging, The Netherlands, http://svi.nl) to remove background noise and increase resolution and signal-to noise ratio. The Classic Maximum Likelihood Estimation (CMLE) algorithm with theoretical PSF was performed for up to 60 iterations. The signal-to-noise ratio for all channels was set between 10 and 30, depending on the individual construct; all other settings were default. Deconvolved images are not scaled to intensity where displayed. 2 μm max projections were made where indicated to account for the approximate volume of a fragmented mitochondrion. For “Proportion of cells with NEMO recruitment” quantification, each cell was cropped from its respective field, max projected, and blinded with other cells in the experimental set. Blinded cells were scored as demonstrating NEMO recruitment or not. For most “% Mitochondria with [protein] recruited” analysis, deconvolved, max-projected images were trained with Ilastik software (Berg et al. 2019) and binary images were generated for the respective particles (Mito, NEMO, etc). FIJI/ImageJ (Schindelin et al. 2012) was used to determine overlap between particles, and percentages were calculated from the fraction of NEMO or OPTN particles overlapping mitochondria to total mitochondria. GABARAPs-positive and LC3B-positive mitochondria were identified by hand due to the low signal-to-noise ratio for the antibodies. For quantification of intensity over time, events were chosen to quantify if they remained within the z plane for most of the sequence. ROIs were manually drawn around the fragmented mitochondrion for each time frame. Intensities for NEMO and/or others were measured for each ROI. Five-frame averages were taken to smooth intensity measurements, and background was calculated from the mean intensity in the respective channel for the first 10 frames and subtracted from every time point. Finally, each intensity measurement was normalized to the maximal intensity recorded for that channel for that event to produce “% of Max Intensity”. For immunoblots, ImageStudio Software (Version 5, Li-Cor) was used to scan bands to ensure no patches were overexposed. ImageStudio was used to subtract background and quantify band intensities as described. Values were graphed and analyzed in GraphPad (Version 9, Prism), except for Euler plots, which were created using RStudio. Images were assembled in Illustrator (Adobe)

## Supporting information

SuppMov1

SuppMov2

SuppMov3

## Acknowledgements

We thank Mariko Tokito for reengineering multiple experimental constructs with various tags and the entire Holzbaur group for invaluable discussion. We gratefully acknowledge National Institute of Neurological Disorders and Stroke (NINDS) (grant NS060698) and the joint efforts of The Michael J. Fox Foundation for Parkinson’s Research (MJFF) and the Aligning Science Across Parkinson’s (ASAP) initiative. MJFF administers the grant ASAP-000350 on behalf of ASAP and itself.

---

**Supplementary Figure 1.**
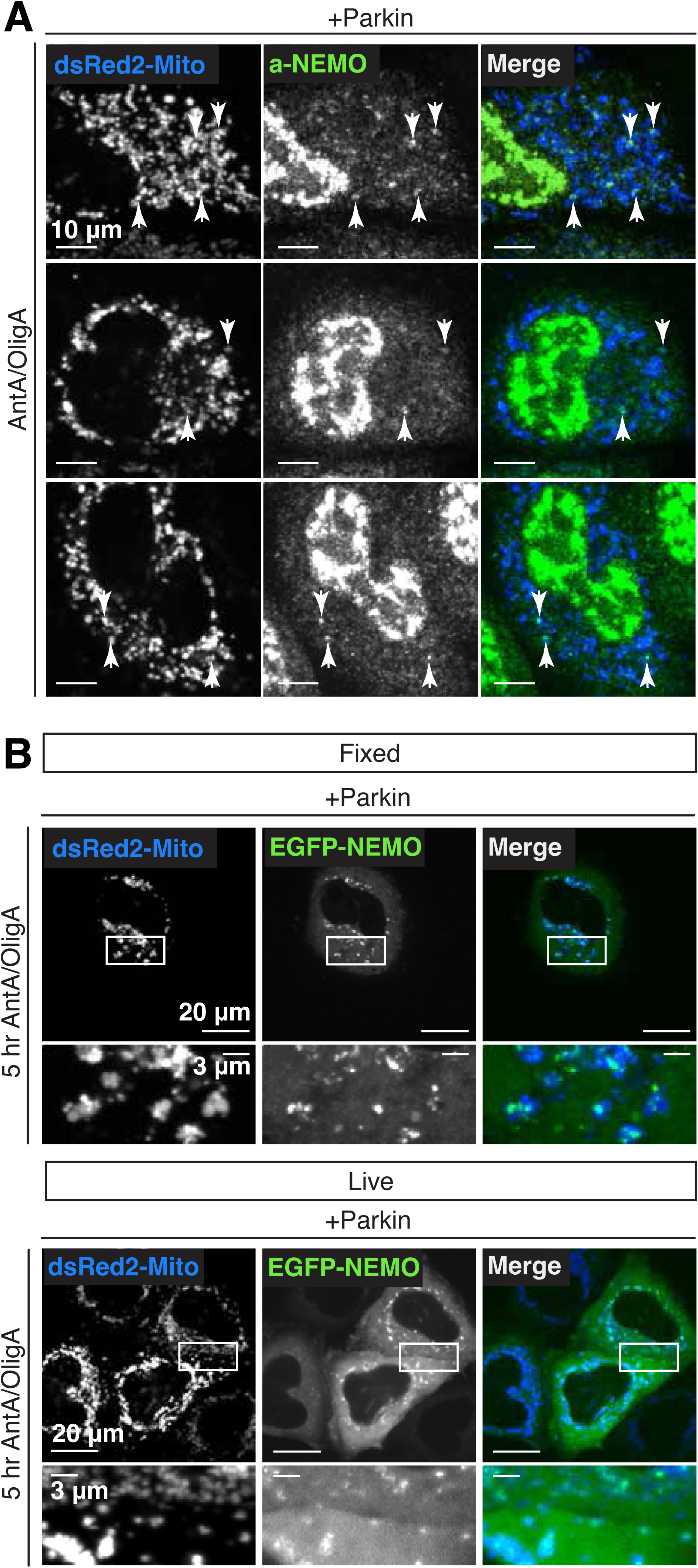
Endogenous NEMO is recruited to depolarized mitochondria and NEMO localization persists up to 5 hr after depolarization. (A) Images of three HeLa cells from two independent experiments expressing dsRed2-Mito and untagged Parkin, fixed after AntA/OligA treatment and labeled with an antibody to NEMO. Arrows indicate mitochondria with NEMO recruitment. Images are max-projected 2 μm. (B) Images of fixed (top) or live (bottom) HeLa cells expressing EGFP-NEMO, dsRed2-Mito, and untagged Parkin, treated with AntA/OligA for 5 hr. Images are max-projected 2 μm.

**Supplementary Figure 2.**
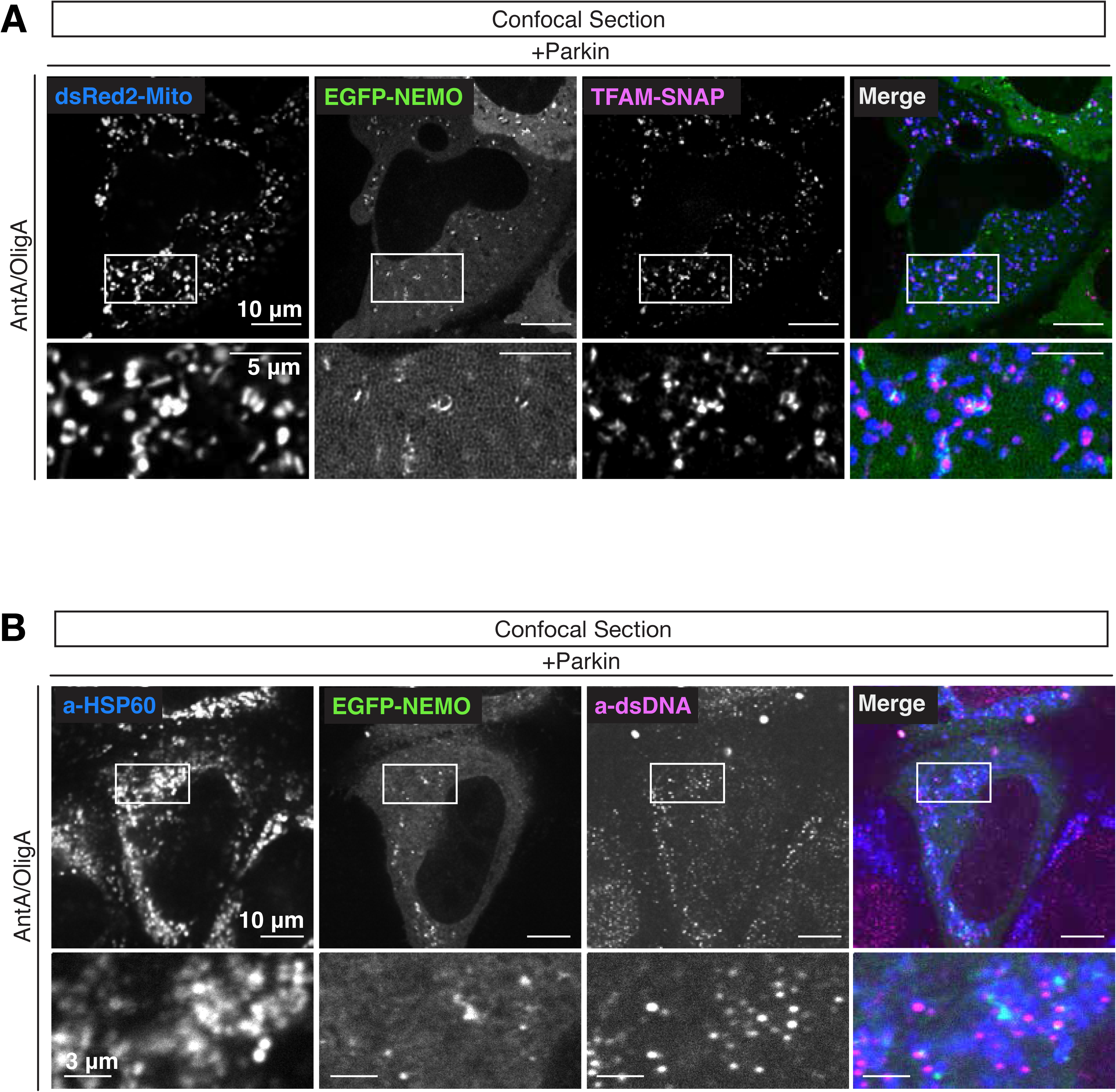
NEMO puncta are not correlated with the localization of mtDNA or mtDNA-associated protein TFAM. (A) Confocal image of live HeLa cells expressing EGFP-NEMO, dsRed2-Mito, SNAP-TFAM, and untagged Parkin, taken after 50-70 min AntA/OligA treatment. (B) Confocal image of HeLa cell expressing EGFP-NEMO and untagged Parkin, fixed after AntA/OligA treatment for 60 min and labeled with antibodies to HSP60 (Mito) and dsDNA.

**Supplementary Figure 3.**
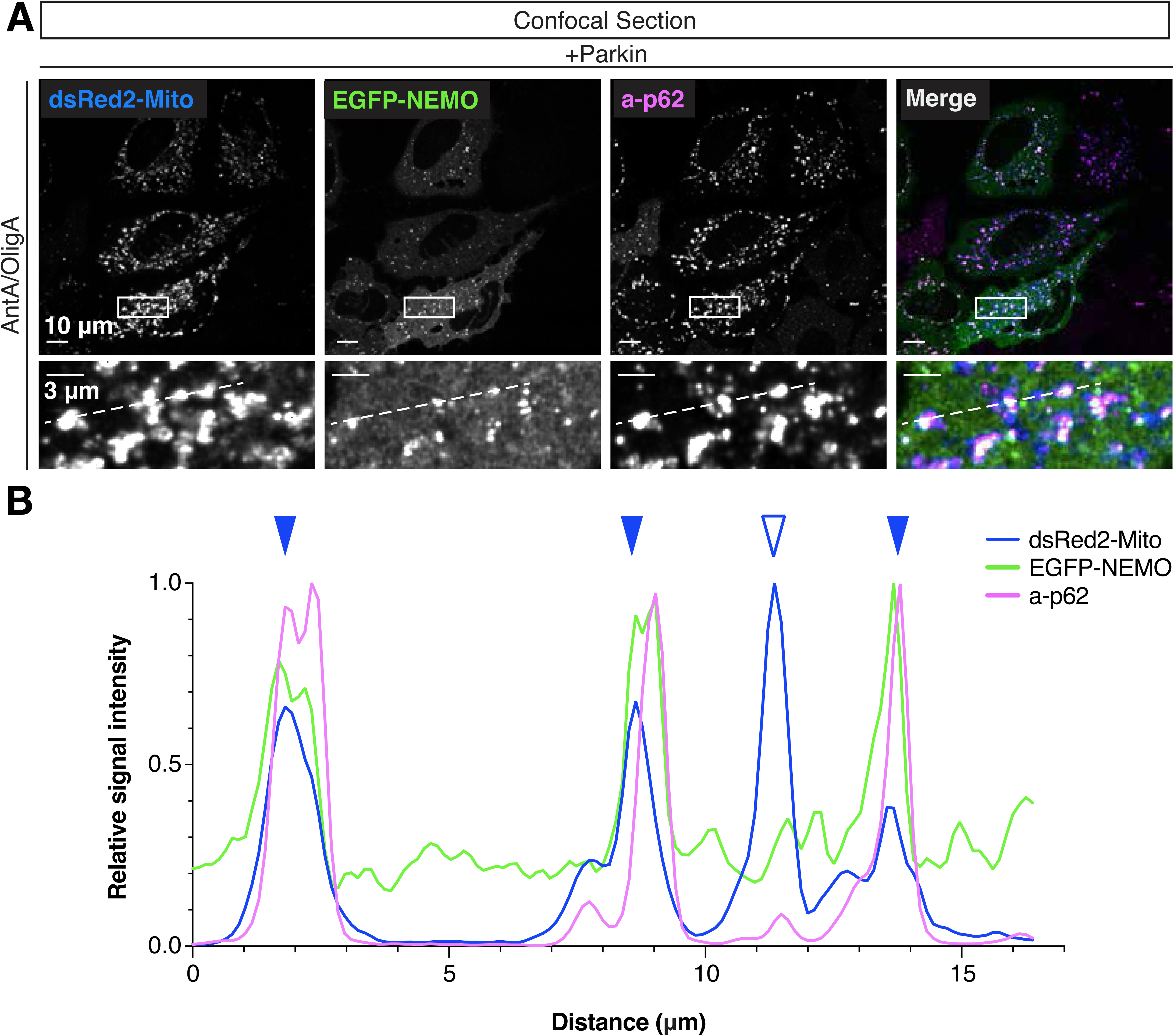
NEMO and endogenous p62 demonstrate extensive colocalization after mitochondrial depolarization. (A) Deconvolved confocal image of HeLa cells expressing EGFP-NEMO, dsRed2-Mito, and untagged Parkin, fixed after 50-70 min AntA/OligA treatment and labeled with an antibody to p62. In zoom images, dotted line indicates the linescan trace in (B). (B) Line scan of relative fluorescence intensities of Mito, NEMO, and p62 in the AntA/OligA-treated cells in (A). Solid arrows indicate mitochondria with recruitment of colocalized NEMO and p62. Unfilled arrow indicates mitochondria with no recruitment of NEMO or p62.

**Supplementary Figure 4.**
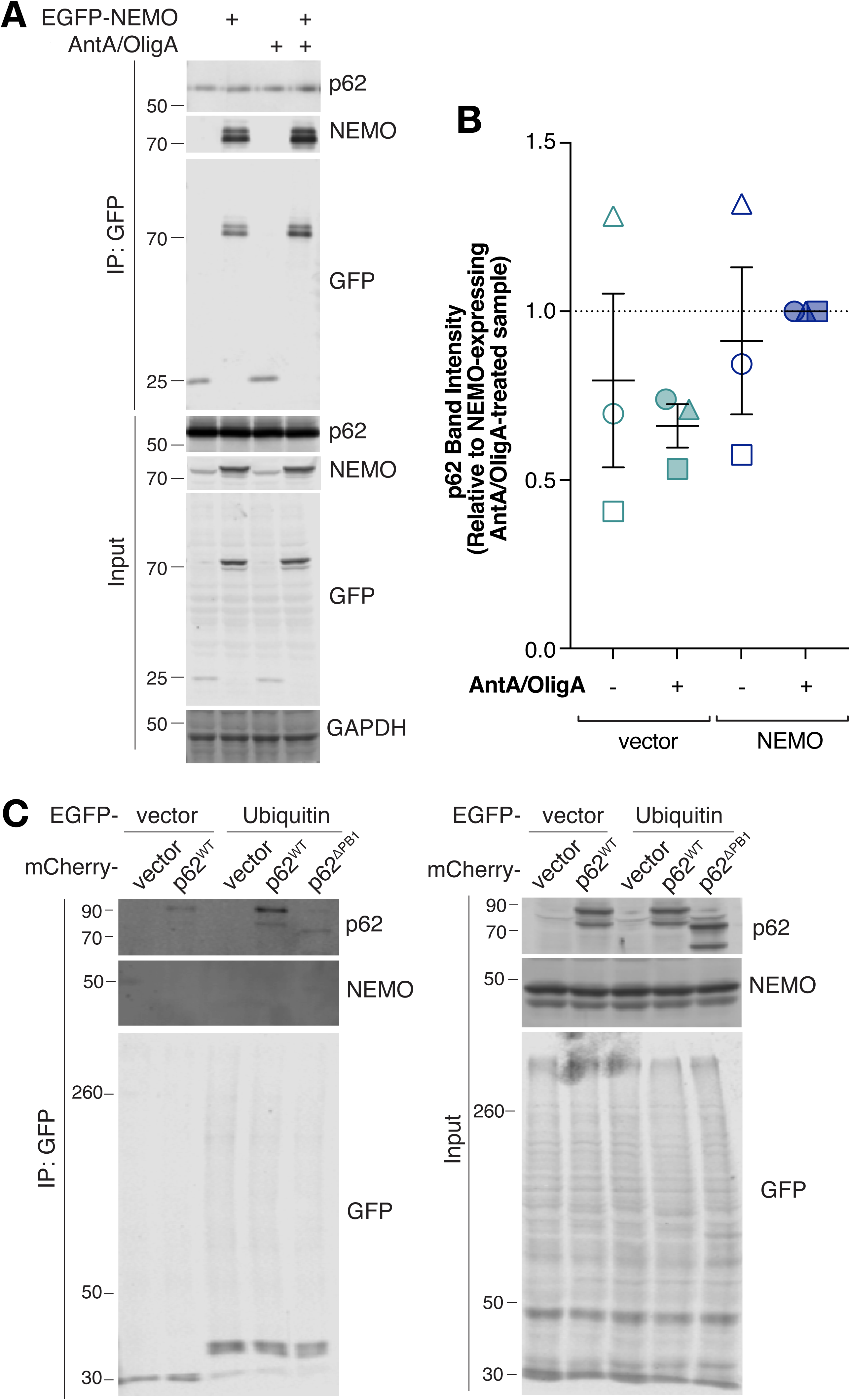
NEMO and p62 are not associated in the cytosolic fraction and p62 is not sufficient to recruit NEMO to ubiquitin. (A) Representative Western blot for immunoprecipitation of EGFP-NEMO (top panels) and input (bottom panels) from HeLa cells expressing Parkin and EGFP-NEMO or EGFP-vector. Cells were treated with 1 hr AntA/OligA where indicated. Numbers on left of images indicate kDa. (B) p62 band intensities in immunoprecipitated fraction (normalized to p62 band in input) relative to intensity in EGFP-NEMO expressing, AntA/OligA-treated samples. Data were collected over three independent experiments. (C) Representative Western blot for immunoprecipitation of EGFP-Ubiquitin or EGFP-vector (left) and input (right) from *p62-/-* cells expressing Parkin, EGFP-NEMO, and mCherry-tagged p62 variant vector. NEMO bands were absent from IP lanes even in high contrast. Numbers on left of images indicate kDa. Experiment performed twice.

**Supplementary Figure 5.**
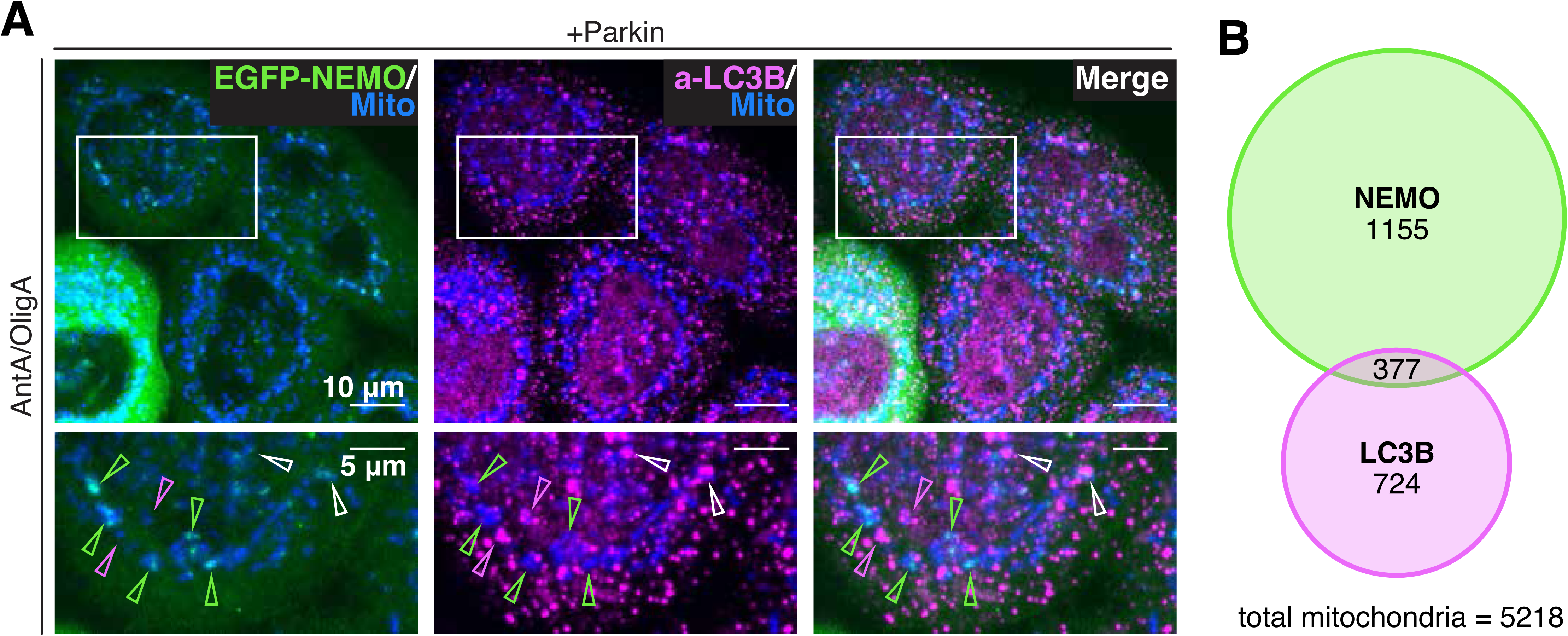
The ATG8 LC3B does not overlap with NEMO on depolarized mitochondria. (A) Image of HeLa cells expressing EGFP-NEMO, dsRed2-Mito, and untagged Parkin, fixed after 60 min AntA/OligA treatment and labeled with an antibody to LC3B. In zoom images (bottom row), arrows indicate mitochondria with only NEMO (green) only LC3B (magenta) or both (white). Images are max-projected 2 μm. (B) Euler diagram displaying the raw number of mitochondria positive for NEMO, LC3B, or both. The remaining mitochondria were positive for neither. Data collected from three independent experiments.

**Supplementary Figure 6.**
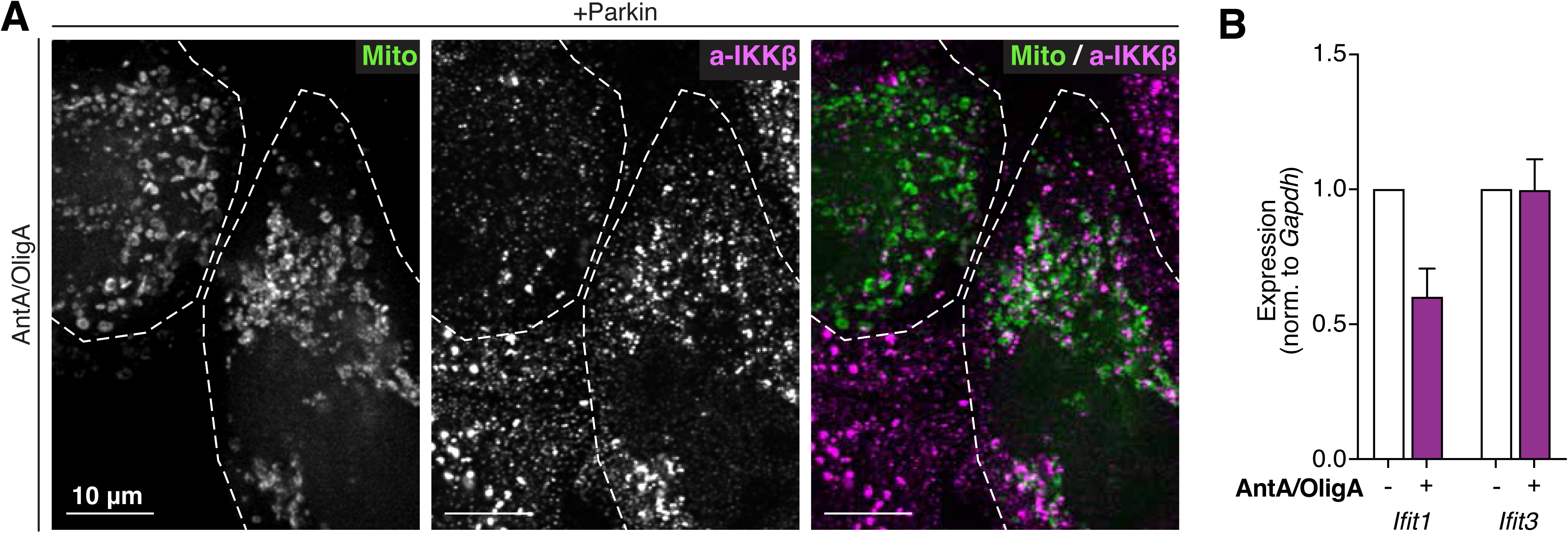
Endogenous IKKβ is recruited to damaged mitochondria and activates NF-κB response in cells. (A) Image of HeLa cells expressing BFP2-Mito (Mito), and untagged Parkin, fixed after 60 min AntA/OligA treatment and labeled with an antibody to IKKβ. Dotted lines indicate cell outlines. Bottom right cell demonstrates extensive IKKβ recruitment to mitochondria while top left cell does not. Images are max-projected 2 μm. (B) RT-qPCR results for HeLa cells expressing Parkin and EGFP-NEMO treated with vehicle (white bars) or AntA/OligA (purple bars). Transcripts for *Ifit1* and *Ifit3* were measured relative to those for *Gapdh*. Data collected over three independent experiments.

Supplementary Movie 1. NEMO is recruited to depolarized mitochondria within 1 hr of AntA/OligA depolarization. Timelapse images of HeLa cells expressing, dsRed2-Mito (Mito), EGFP-NEMO, and Parkin (untagged), imaged at 1 frame per min in a confocal section. Time (hour:min) beginning at frame 6 indicates addition of AntA/OligA.

Supplementary Movie 2. OPTN and NEMO are recruited to depolarized mitochondria with similar kinetics. Timelapse images of HeLa cells expressing (from bottom left in clockwise order) Halo-OPTN, dsRed2-Mito (Mito), EGFP-NEMO, and Parkin (untagged), imaged at 1 frame per min in a confocal section. Time (hour:min) beginning at frame 6 indicates addition of AntA/OligA.

Supplementary Movie 3. p62 and NEMO recruitment to depolarized mitochondria is correlated in space and time. Timelapse images of HeLa cells expressing (from bottom left in clockwise order) mCherry-p62, sBFP2-Mito (Mito), EGFP-NEMO, and Parkin (untagged), imaged at 1 frame per min in a confocal section. Time (hour:min) beginning at frame 7 indicates addition of AntA/OligA.

